# A novel preparation of the entire isolated CNS unveils the modulatory influences of supra-pontine structures on brainstem and spinal networks

**DOI:** 10.1101/2022.06.24.497519

**Authors:** Atiyeh Mohammadshirazi, Rosamaria Apicella, Giuliano Taccola

## Abstract

Several spinal motor output and essential rhythmic behaviors are controlled by supraspinal structures. Although of great advantage in deciphering their functional organization, existing preparations of the isolated brainstem and spinal networks only focus on local circuitry. The goal of our study has been to better characterize the contribution of higher centers to the neuronal networks involved in respiration and locomotion. Thus, a novel in vitro preparation from the isolated CNS of neonatal rodents was introduced to simultaneously record a stable respiratory rhythm from both cervical and lumbar ventral roots (VRs).

Selective electrical pulses supplied to the pons and medulla evoked distinct VR responses in rostrocaudal direction with a staggered onset, while stimulation of ventrolateral medulla resulted in higher events from homolateral VRs. Moreover, electrical stimulation of a lumbar dorsal root (DR) elicited responses even from cervical VRs, albeit small and delayed, confirming functional ascending pathways. Furthermore, prototypical fictive locomotion was induced by trains of pulses applied to either the ventrolateral medulla or a DR.

By progressively removing higher centers, duration of respiratory burst was reduced after a precollicular decerebration, which also affected the area of lumbar DR and VR potentials elicited by DR stimulation while frequency of respiration increased after a following pontobulbar transection. Keeping legs attached to the CNS allows for expressing the respiratory rhythm during peripheral stimulation of limbs.

The study demonstrates that supra-pontine centers regulate the spontaneous respiratory rhythm, as well as electrically-evoked reflexes and spinal network activity. Thus, the current approach contributes to clarifying the modulatory influence of the brain on the brainstem and spinal micro-circuits.

## METHODS

### In vitro preparation of the isolated entire CNS

All procedures were approved by the International School for Advanced Studies (SISSA) ethics committee and are in accordance with the guidelines of the National Institutes of Health (NIH) and with the Italian Animal Welfare Act 24/3/2014 n. 26, implementing the European Union directive on animal experimentation (2010/63/EU). All efforts were made to minimize the number and suffering of animals. A total of 140 postnatal Wistar rats (P0-P4) of random sexes were used.

As graphically summarized by the timeline in Fig. 1A, 7-11 minutes of cryoanesthesia (Danneman and Mandrell, 1997; Phifer and Terry, 1986; Zimmer et al., 2020) anticipates surgical procedures. After disappearance of tail pinch reflex, the forehead was ablated at the level of the orbital line, the skin removed from the animal’s skull and back, and the chest and forelimbs ventrally detached. The preparation was then placed on a Sylgard-filled petri dish under a microscope and fully covered with oxygenated Krebs solution, which was frequently replaced. Krebs solution contained (in mM): 113 NaCl, 4.5 KCl, 1 MgCl_2_7H_2_O, 2 CaCl_2_, 1 NaH_2_PO_4_, 25 NaHCO_3_, and 30 glucose, gassed with 95% O_2_-5% CO_2_, pH 7.4, 298 mOsm/kg. Afterwards, craniotomy and ventral and dorsal laminectomies were performed to expose the entire CNS, which was then isolated from the olfactory bulbs down to the cauda equina by carefully transecting all cranial nerves, dorsal roots (DRs) and ventral roots (VRs; Nicholls et al., 1990). Fig. 1B pictures the whole CNS preparation from a P1 newborn in dorsal and ventral views. On average, dissection procedures lasted about 30 minutes. A post-dissection resting period of 15 minutes was systematically respected after surgical dissection (Fig. 1A).

**Figure 1.**
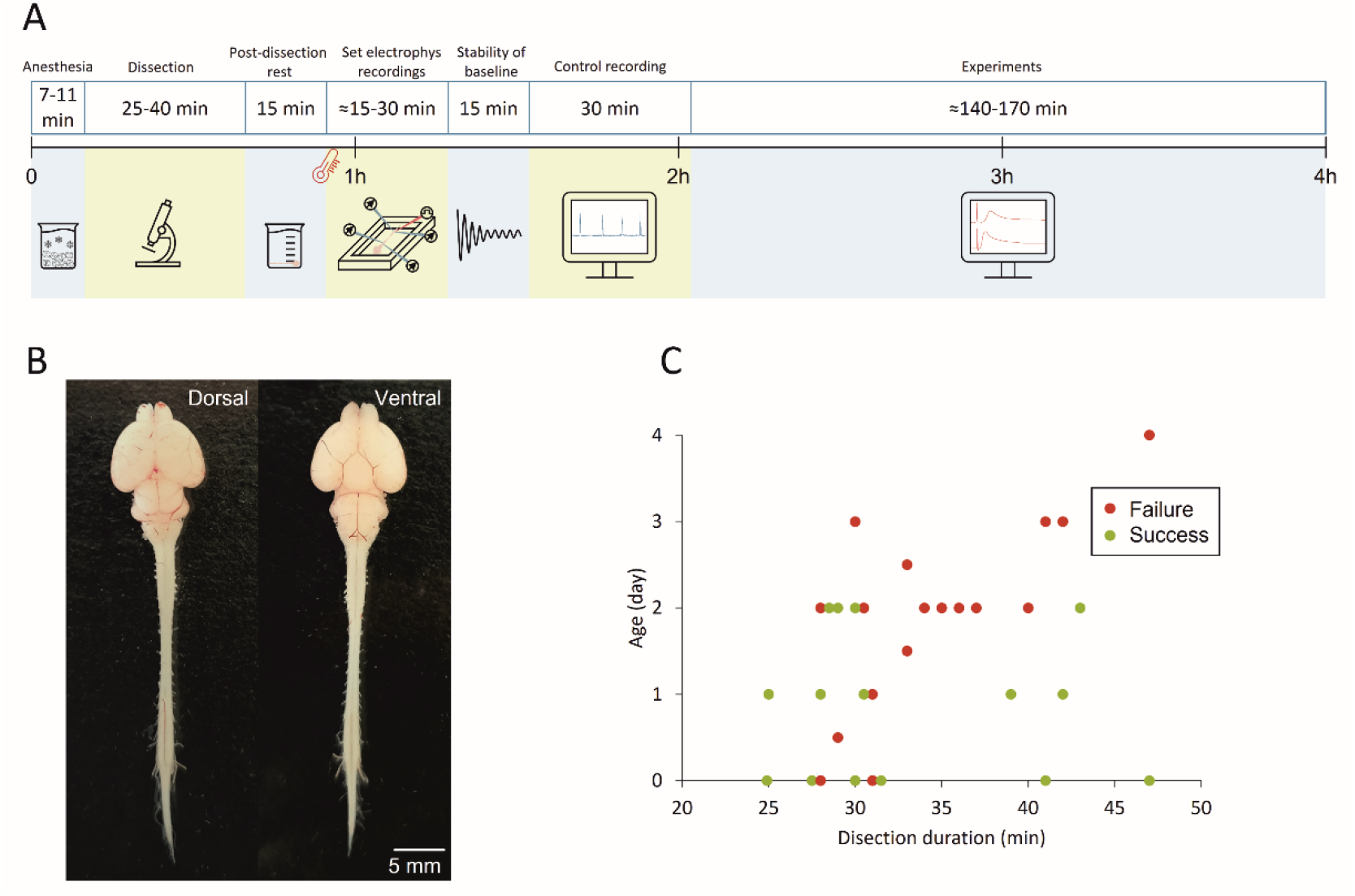
Experimental design of the study. (A) A timeline describing the experimental procedures, from cryoanesthesia to dissection, post-dissection rest, setting of electrophysiological recordings, pause to stabilize the baseline, and eventually control and experimental recordings. Note that as soon as the preparation was placed in the recording chamber, bath temperature was progressively raised to 27°C. (B) Photos displaying dorsal and ventral views of the isolated entire CNS preparation from a P1 newborn. (C) Scatter plot describes the duration of fictive respiration for each preparation arranged by age and time of dissection. Green dots correspond to successes (>4 hrs of fictive respiration), while red dots represent failures (>4 hrs of fictive respiration).

For electrophysiological recordings, the preparation was then placed ventral side up in a recording chamber (Fig. 1A) continuously perfused with Krebs solution (7 mL/min), while the bath temperature was progressively raised and maintained in the range of 25 to 27°C by a single channel temperature controller (TC-324C Warner Instruments, USA). Once the baseline stabilized (15 min; Fig. 1A), 30 mins of stable control (Fig. 1A) were acquired through extracellular recordings.

For the preparation of the entire CNS with legs attached, dorsal and ventral laminectomies were performed down to the lowest thoracic level (Th13), while preserving lumbo-sacral vertebrae and nerves attached to hindlimbs. DRs and VRs were kept as long as possible, removing only dorsal root ganglia (DRG).

Plots in Figs. 1C and 9B, report age of the animal on the x-axis and length of surgical dissection on y-axis, identifies a bottom-left region (younger preparations undergoing fast surgical procedures) where preparations showed the highest percentage of successfully recorded respiratory rhythms for at least 4 hours from the anesthesia at the beginning of tissue isolation. Based on the outcome of this set of preliminary experiments, P0-2 newborns were selected for the rest of the study.

### Extracellular recordings

DC-coupled recordings were acquired from VRs and DRs through tight-fitting suction electrodes connected to a differential amplifier (DP-304, Warner Instruments, Hamden, CT, USA; high-pass filter – 0.1 Hz, low-pass filter = 10 kHz, gain X 1000), then digitized (Digidata 1440, Molecular Devices Corporation, Downingtown, PA, USA; digital Bessel low-pass filter at 10 Hz; sampling rate = 50 kHz) and visualized real-time with the software Clampex 10.7 (Molecular Devices Corporation, Downingtown, PA, USA).

### Electrical Stimulation

Single and repetitive rectangular electrical pulses (duration = 0.1 ms, frequency = 0.03 Hz) were delivered to caudal DRs (L4-S1) through a programmable stimulator (STG4002, Multichannel System, Reutlingen, Germany) using bipolar glass suction electrodes with two close silver wires (500-300 µm). Intensity of stimulation (6-800 µA) is expressed as times to threshold, where the latter is the lowest intensity supplied to a DR to elicit an appreciable depolarizing potential from the homolateral VR. As for input-output experiments, to elicit motor potentials in response to DR stimulation (DRVRPs), 30 single pulses (duration = 0.1 ms) at intensities of 1, 1.5, 2, 3, 5 x Thr were delivered at a frequency of 0.03 Hz.

Punctiform stimulation of multiple sites of the brain was conducted with a custom-made bipolar concentric electrode composed of an internal 250 µm width stainless steel electrode (UE KK1, FHC, Bowdoinham, USA) and a helical silver wire wrapped around the tip of a glass pipette (700 µm diameter tip). Multisite stimulation of the brainstem consists in a train of 30 rectangular pulses at 0.03 Hz (duration = 1 ms; intensity = 2 x Thr, 1800-5000 µA). In experiments involving brainstem stimulation, threshold was defined as the lowest intensity applied to the ventrolateral medulla to obtain an appreciable depolarizing response from lower lumbar VRs. In each experiment, 10 distinct stimulating spots (named A-J) were consistently found on each side of the brainstem based on the anatomy of visible ventral arteries. Two stimulating sites are positioned on the median basal artery (namely A on the most rostral pons, and G at the intersection of basilar and anterior inferior cerebellar arteries). B is positioned about 0.5 mm laterally from A. The spots named C, D and E are aligned on the pons, equally interspaced by 0.5 mm and span from the median basal artery (C) to the lateral extremity of the ventral brainstem (E). On rostral caudal direction, D is equidistant between superior and inferior cerebellar arteries. The F stimulating spot is placed on the anterior inferior cerebellar artery between the median basal artery and lateral ventral edge of the brainstem. H and I are aligned on the medulla, proximal (H) and distal (I) to the median basal artery. On rostral caudal direction, they are equidistant from the inferior cerebellar artery and the first cervical VR. J is located on the first cervical segment of the spinal cord, laterally to the ventral spinal artery.

In a subgroup of preparations, the stimulation site was visually confirmed at the end of the experiment by electrolytically destroying the area of stimulation through strong electrical pulses (intensity = 16 mA, duration = 5 ms) delivered on the surface of pyramids (spot H) in the ventrolateral medulla.

Fictive locomotion (FL) patterns were recorded from the left (l) and right (r) L2 VRs (flexor motor commands) and from l and r L5 VRs (extensor motor commands). Alternating discharges between homolateral L2 and L5 VRs and between homosegmental VRs is considered the distinct feature of FL (Kiehn, 2006). FL was electrically evoked by trains of rectangular single pulses applied to DRrL6-S1 (160 single pulses at 2Hz, pulse duration = 0.1 ms, intensity = 15-37.5 µA or 1.5-3.5 Thr) or to the pyramid in the ventrolateral medulla (trains of single pulses at 1-2 Hz, pulse duration = 1-5 ms, intensity = 0.5-4.5 mA).

### Serial transection experiments

In a subgroup of experiments, supra-pontine structures were ablated from the whole CNS preparation by two serial horizontal transections. Firstly, precollicular decerebration was performed by surgically cutting the brain rostral to the fifth cranial nerves at the level of superior cerebellar arteries and caudal edge of inferior colliculi (Voituron et al., 2005). One hour later, a second transection was carried out at the level of the ninth cranial nerves to separate pons and medulla. Before each cut, suction electrodes on cervical VRs were released to avoid any nerve damage and a new suction was adopted after cutting. In experiments with DR train stimulation, a further midthoracic transection (at the level of thoracic 4/5) was adopted to compare, in the same animal, FL in the intact CNS vs. the isolated spinal cord. Since in these experiments, midthoracic transection did not affect stability of lumbar VR signals, suctions were not released, thus allowing a direct comparison between the amplitude of signals before and after spinal transection.

### Data analysis

To remove electrical interference, original traces were notched at 50 Hz through Clampfit 11.2 software (Molecular Devices Corporation, PA, USA). All spontaneous rhythmic motor discharges displaying large amplitude depolarizations synchronous among bilateral VRs and appearing at regular intervals, are ascribed to respiratory bursts. A burst is defined as a period of sustained membrane depolarization that originates with a rapid onset from the baseline and remains above a preset threshold (usually five times the standard deviation of baseline noise; Bracci et al., 1996). The time during which the membrane potential remains above the preset threshold is defined as burst duration (Bracci et al., 1996). Rhythmic discharges were also characterized based on their period, defined as the time between peaks of two consecutive cycles (Dose et al., 2014). The ratio between standard deviation and mean value provide the coefficient of variation (CV), which is an index of consistency of responses (Taccola et al., 2020).

Averaged electrically evoked reflex responses were obtained by averaging at least 15 traces not corrupted by any spontaneous activity.

Conduction velocity is calculated by dividing the time to peak of each response by the distance between stimulating and recording sites, as measured by a microcalibrated dial caliper (sensitivity = 20 µm).

Phase coupling between pairs of VRs was ascertained by the correlation coefficient function (CCF) using Clampfit 11.2 software. A positive CCF value ≥ 0.5 states that rhythmic signals from two VRs are synchronous while CCF ≤ -0.5 accounts for alternating patterns (Dose et al., 2014).

### Statistical analysis

Statistical analysis was performed with GraphPad InStat 3.10 (Inc., San Diego, California, USA). All data in boxplots show sample median (horizontal segment), 75^th^ and 25^th^ percentiles (top and bottom edges of box) and 1.5 times the interquartile range (whiskers). Number of animals is indicated as n in the Results, and data are reported as mean ± SD values. Before assessing statistical differences among groups, a normality test was performed to select the use of either parametric or non-parametric tests.

Accordingly, parametric data were analyzed with paired or unpaired t-test, one-way analysis of variance (ANOVA) and repeated measure ANOVA; whereas Mann-Whitney Test, Kruskal-Wallis Test, and Wilcoxon matched-pairs signed-ranks test are used for non-parametric data. Multiple comparisons ANOVA was followed by Tukey-Kramer Multiple Comparisons Test, Fisher LSD, or Dunn’s Method. Differences were considered statistically significant when P ≤ 0.05.

## RESULTS

### The entire CNS in vitro reveals a spontaneous and stable fictive respiratory rhythm

The expression of spontaneous respiratory motor patterns is a sign of the functionality of neuronal networks from the brainstem-spinal cord *in vitro* (Smith et al., 1990). Likewise, in a sample preparation of the entire CNS isolated from newborn rats, we recorded respiratory-related rhythmic discharges at a frequency of 0.04 Hz, which appeared synchronous (CCF = 0.84) among cervical and lumbar ventral roots (Fig. 2A). The average burst is reported in Fig. 2B, showing a duration of 1.47 s for VRrC1 and 1.09 for VRrL5 and a peak amplitude equal to 0.25 mV and 0.06 mV respectively. Interestingly, double bursts only seldom appeared, as visualized in the magnification of Fig. 2C where double- (red star) and single-peaked respiratory events follow in a row.

**Figure 2.**
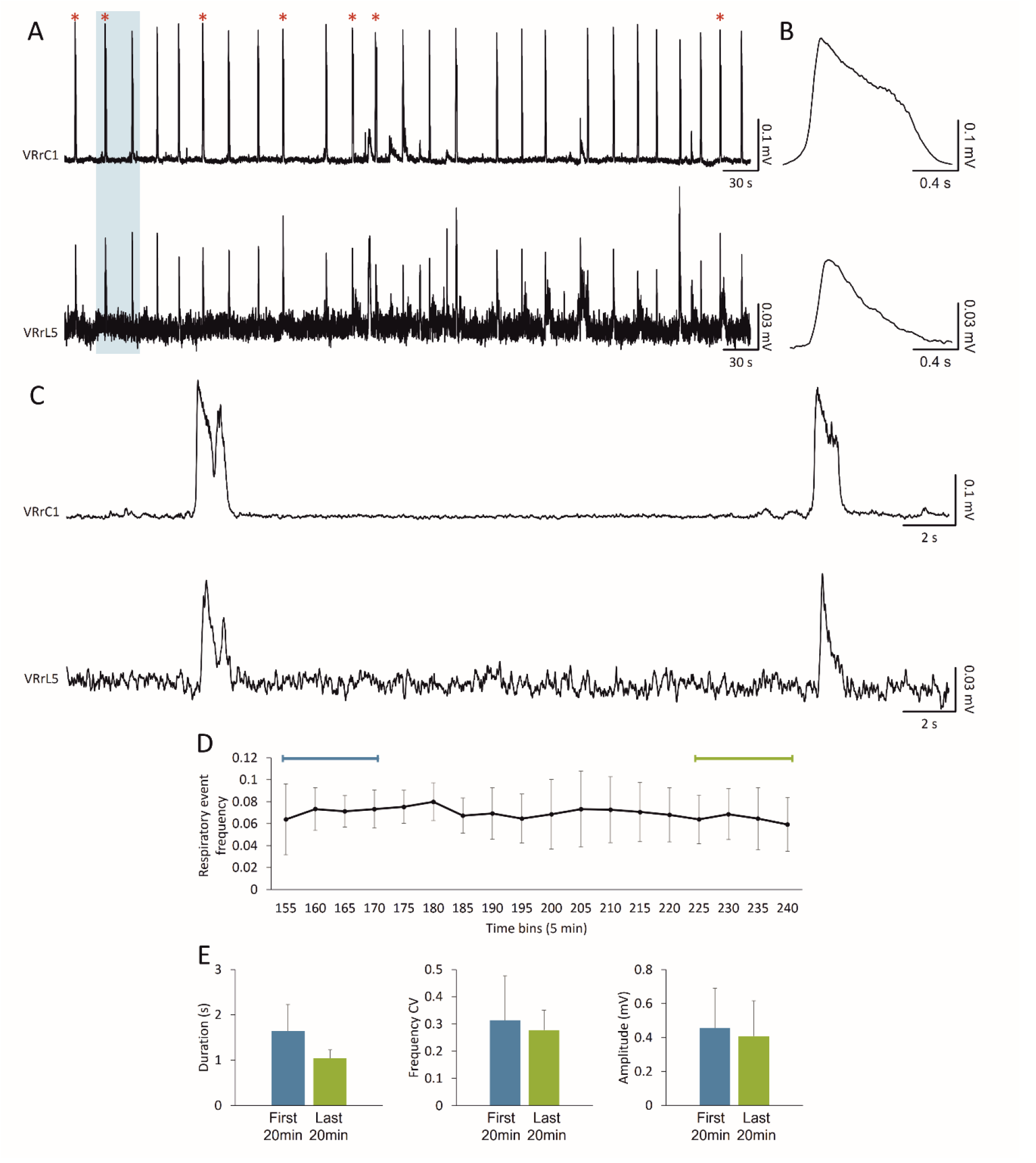
In vitro preparation of the entire CNS expresses a stable fictive respiration for four hours. (A) Spontaneous and synchronous respiratory bursts are recorded for up to four hours from both cervical and lumbar VRs. The occurrence of sporadic double bursts is tagged by red stars. (B) Average bursts summarize shape and duration of cervical and lumbar respiratory events. (C) Magnification of traces in the shaded rectangle in A. A double and a single peak burst follow one another on both cervical and lumbar VRs. (D) The average time course shows the frequency of fictive respiration for the entire length of the experiment for bin = 5 min. The time indicated lapses from the induction of anesthesia. (E) Respiratory events in the first and last 20 minutes of the time course in D remain unchanged as for duration, regularity of rhythm expressed as frequency CV, and amplitude.

Pooled data from twelve experiments indicate that, in the first 30 minutes of continuous recordings, the spontaneous respiratory rhythm has a frequency of 0.06 ± 0.03 Hz, with single bursts that last on average 1.80 ± 0.52 s and peak amplitude of 0.24 ± 0.13 mV. In six of those preparations, fictive respiratory events seldom appeared double-peaked (in average, 8.17 ± 2.48 double-peaked burst out of 102.17 ± 63.02 total respiratory events) and eventually turned into single bursts in the following 30 mins. However, the sporadic occurrence of double bursts did not affect mean rhythm frequency (P = 0.448; unpaired t-test), burst duration (P = 0.449; unpaired t-test), nor peak amplitude (P = 0.240; Mann-Whitney Test).

To prove stability of the fictive respiratory rhythm from the entire CNS preparation in vitro, long recordings were continuously performed for at least 4 hours right after tissue isolation. Fig. 2D traces the time course of the mean rhythm frequency in 5-minute bins for five experiments. Respiratory events recorded in the first 20 minutes were similar to the ones recorded at the end of the time course as for frequency (P = 0.536; paired t-test), duration (P = 0.050; paired t-test), frequency CV (P = 0.680; paired t-test) and amplitude (P = 0.386; paired t-test; Fig. 2E, Table 1).

**Table 1.** Features of fictive respiratory events at the onset and the end of electrophysiological experiments.

|  | First 20 min | Last 20 min |
| --- | --- | --- |
| Frequency | 0.07 ± 0.03 | 0.07 ± 0.02 |
| Duration (s) | 1.65 ± 0.59 | 1.04 ± 0.19 |
| Frequency CV | 0.31 ± 0.16 | 0.28 ± 0.07 |
| Amplitude (mV) | 0.46 ± 0.24 | 0.41 ± 0.21 |

The stable fictive respiratory rhythm recorded for more than four hours demonstrates that the entire CNS in vitro is a sound preparation for studying the respiratory network in a more intact experimental setting.

### Punctiform electrical stimulation of the ventral surface of the brainstem and pons elicits distinct motor responses along lumbar segments

To assess descending input conduction along spinal segments in the CNS preparation, motor evoked potentials were induced by serial pulses of electrical stimulation on different sites of the ventral brainstem. On each preparation, electrical pulses were delivered to ten loci on each side of the brainstem, ranging from higher pontine structures to the upper cervical cord (locations are detailed in the method section).

The cartoon in Fig. 3A identifies the stimulating sites using different letters and a color code. The responses below were serially evoked, in the same preparation, by electrical stimulation of distinct sites and were recorded from the homolateral VRrL5. In a sample experiment, the stimulating electrode was moved in a cranial-rostral direction, generating larger and more delayed responses (Fig. 3A). Medial rostral pons (indicated by the letter “A” in Fig. 3A) were the highest sites of stimulation that still generated appreciable evoked potentials from lumbar motor pools (L5), with a peak of 0.04 mV and a time to peak of 0.14 s. The largest (Fig. 3B) and earliest (Fig. 3C) motor evoked response, with a peak of 1.84 mV and a time to peak of 0.08 s, was obtained by pulses supplied to the ventrolateral surface of the first cervical spinal segment (named as J). On the other hand, the most efficient brainstem stimulation corresponded to the supply of pulses to the pyramid (as indicated by the letter “H”) with responses of 1.68 mV amplitude and 0.09 s time to peak. Contrariwise, stimulation of lateral rostral pons and higher brain structures (dorsal cerebellum, and both ventral and dorsal mesencephalon and cortex) always failed to elicit any reflexes from VRs (data not shown).

**Figure 3.**
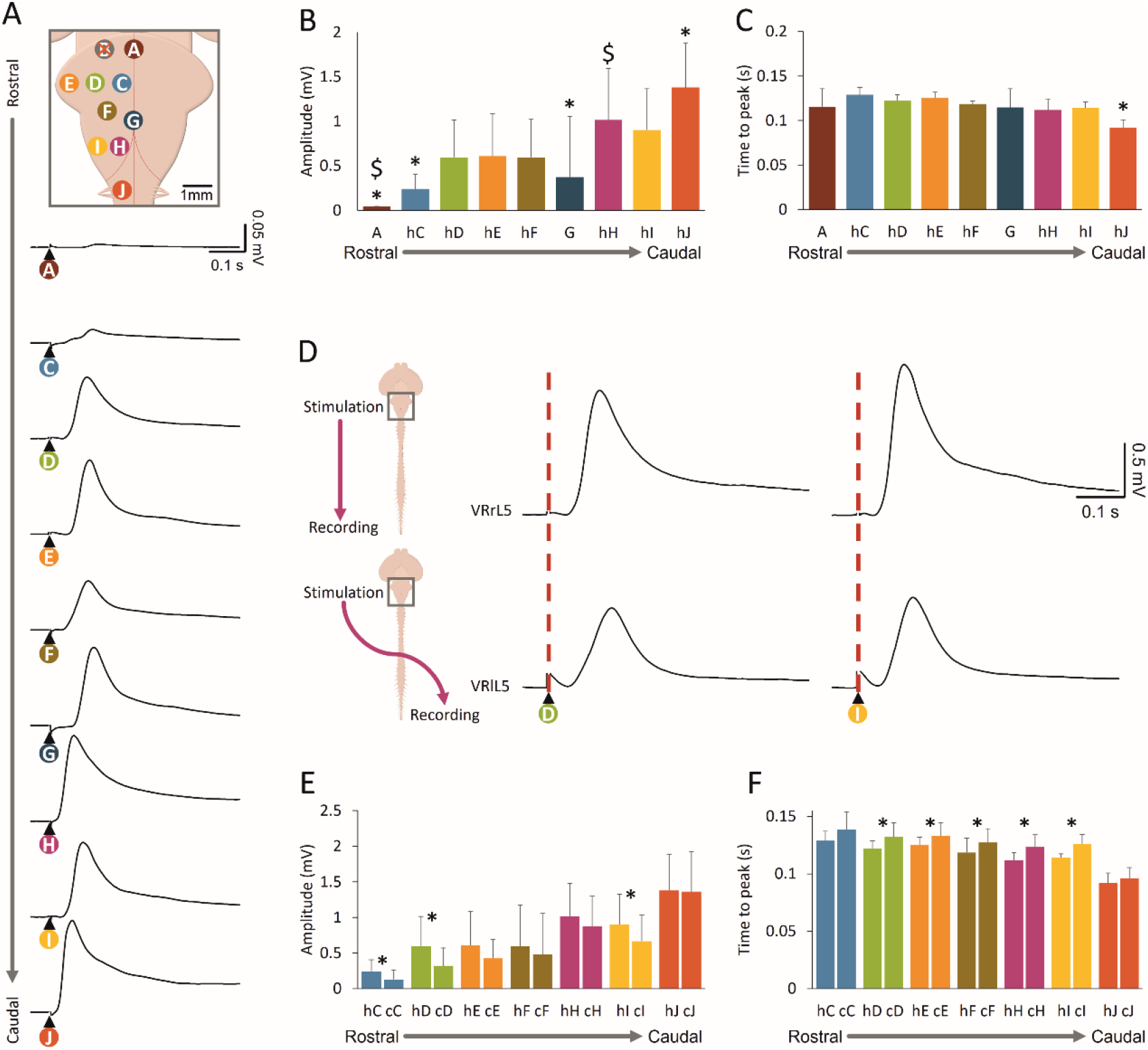
Distinct motor responses from lumbar VRs are evoked by punctiform electrical stimulation of the ventral surface of the brainstem and pons. (A) Different sites of pons and medulla were serially stimulated by a custom-made punctiform electrode (intensity = 2 mA, duration = 1 ms) and mean motor potentials were recorded from VRrL5. Each site of stimulation is indicated with a different color and letter in the schematic cartoon, which is calibrated to the real sample dimensions. Please note that only one side of the cartoon is labeled. Serial stimulation of distinct sites evokes motor responses of different amplitude (B, *P < 0.001) and latency (C, *P < 0.001). (D) Motor responses are elicited from r-l VRL5 by serially stimulating “D” (left panel) or “I” spots (right panel) on the right medulla (same experiment as in A). For each pulse, homolateral (upper) and contralateral (bottom) responses are simultaneously acquired. Average traces come from 15 superimposed sweeps. (**E**) Histogram reports the peak amplitude of homolateral and contralateral responses by serially stimulating different sites. Homolateral responses evoked by stimulation of “C” (*P = 0.006), “D” (*P = 0.010), and “I” (*P = 0.023) are significantly higher than contralateral ones. (**F**) Histogram summarizes the time to peak of homolateral and contralateral responses when serially stimulating different sites. Homolateral responses to pulses applied to “D” (*P = 0.043), “E” (*P = 0.031), “F” (*P = 0.020), “H” (*P < 0.001), and “I” (*P = 0.018) are significantly faster in comparison to contralateral.

Average data pooled from 5-8 experiments describes responses evoked by serial multi-site electrical stimulation in terms of amplitude (Fig. 3B) and time to peak (Fig. 3C) and reports significantly larger responses from cervical spinal stimulation compared to rostral pulses (Fig. 3B; “A”, “C” and “G”; P < 0.001; Kruskal-Wallis Test). Moreover, compared to all homolateral responses, the one originating from stimulation of the ″J″ site appears sooner (Fig. 3C; P < 0.0001; ANOVA test followed by Tukey-Kramer Multiple Comparisons Test).

The most rostral impulses that generated VRPs in three out of six experiments were delivered to the medial rostral pons (as indicated by the letter ″A″) still generating responses that were significantly lower compared to the ones elicited by pyramid stimulation (P < 0.001; Kruskal-Wallis Test).

To explore whether homolateral and contralateral stimulation evoked different spinal lumbar responses, simultaneous recordings were acquired from the right and left VRL5 when electrical pulses were serially applied to multiple sites of the brainstem-upper cervical cord. Fig. 3D show simultaneous recordings from bilateral VRL5 during electrical stimulation of rostro-medial pons (“D”) and ventro-lateral (“I”) medulla. On both sites, responses were higher and appeared earlier for homolateral (peak_D_ = 1.20 mV, time to peak_D_ = 0.11 s; peak_I_ = 1.45 mV, time to peak_I_ = 0.11 s) vs. contralateral stimulation (peak_D_ = 0.77 mV, time to peak_D_ = 0.13 s; peak_I_ = 0.88 mV, time to peak_I_ = 0.11 s). In 5-8 experiments, the majority of stimulating configurations generated homolateral (h) and contralateral (c) evoked responses of unchanged amplitude, excluding the pons (“C” and “D”) and ventro lateral medulla (“I”; Fig. 3E). As for latency of evoked responses expressed as time to peak, contralateral stimulation elicited delayed VRPs in all cases, except for the site indicated by the letter “C”, which is very close to the brainstem midline. Average values are reported in Table 2.

**Table 2.** Amplitude of homolateral and contralateral VR potentials elicited by punctiform electrical stimulation of different spots on the ventral surface of the brainstem.

| Location | A<br>(n=3) | C<br>(n=6) | D<br>(n=8) | E<br>(n=8) | F<br>(n=7) | G<br>(n=6) | H<br>(n=8) | I<br>(n=7) | J<br>(n=8) |
| --- | --- | --- | --- | --- | --- | --- | --- | --- | --- |
| Homolateral peak amplitude (mV) | 0.04 ± 0.00 | 0.24 ± 0.16 | 0.6 ± 0.48 | 0.61 ± 0.48 | 0.6 ± 0.43 | 0.37 ± 0.68 | 1.02 ± 0.58 | 0.9 ± 0.46 | 1.38 ± 0.5 |
| Homolateral time to peak (m) | 0.12 ± 0.02 | 0.13 ± 0.01 | 0.12 ± 0.01 | 0.13 ± 0.01 | 0.12 ± 0.00 | 0.12 ± 0.02 | 0.11 ± 0.01 | 0.11 ± 0.01 | 0.09 ± 0.01 |
| Contralateral peak amplitude (mV) |  | 0.13 ± 0.13 | 0.32 ± 0.25 | 0.43 ± 0.26 | 0.48 ± 0.37 |  | 0.88 ± 0.58 | 0.67 ± 0.42 | 1.36 ± 0.57 |
| Contralateral time to peak (m) |  | 0.14 ± 0.02 | 0.13 ± 0.01 | 0.13 ± 0.01 | 0.13 ± 0.01 |  | 0.12 ± 0.01 | 0.13 ± 0.01 | 0.1 ± 0.01 |
| P value of homolateral and contralateral amplitude comparison |  | 0.006<br>(paired t-test) | 0.010<br>(paired t-test) | 0.110<br>(Wilcoxon matched-pairs signed-ranks test) | 0.097<br>(paired t-test) |  | 0.210<br>(paired t-test) | 0.023<br>(paired t-test) | 0.894<br>(paired t-test) |
| P value of homolateral and contralateral time to peak comparison |  | 0.143<br>(paired t-test) | 0.043<br>(paired t-test) | 0.031<br>(Wilcoxon matched-pairs signed-ranks test) | 0.020<br>(paired t-test) |  | < 0.001<br>(paired t-test) | 0.018<br>(paired t-test) | 0.123<br>(paired t-test) |

In summary, descending input recorded caudally from bilateral VRs of the lower lumbar cord shows the recruitment of specific functional pathways, having distinct latencies and motoneuronal recruitment mirroring the rostro-caudal supply of brainstem pulses.

To explore the organization of descending spinal pathways recruited by brainstem stimulation, electrical pulses were delivered to the pyramid while responses from homolateral VRs at several spinal levels were used to calculate conduction velocity. In Fig. 4A, evoked motor potentials were acquired from five spinal segments (from C1 to L5) in correspondence to stimulation of the left medulla. Responses became lower and slower when recorded from more caudal spinal segments. While the upper cervical peak appeared sooner (C1; 0.02 s) and was more similar to lower cervical regions (C6; 0.02 s), the thoracic VR response was delayed (T9; 0.04 s) and even more delayed were the responses derived from upper (L1; 0.05 s) and lower (L5; 0.09 s) lumbar segments.

**Figure 4.**
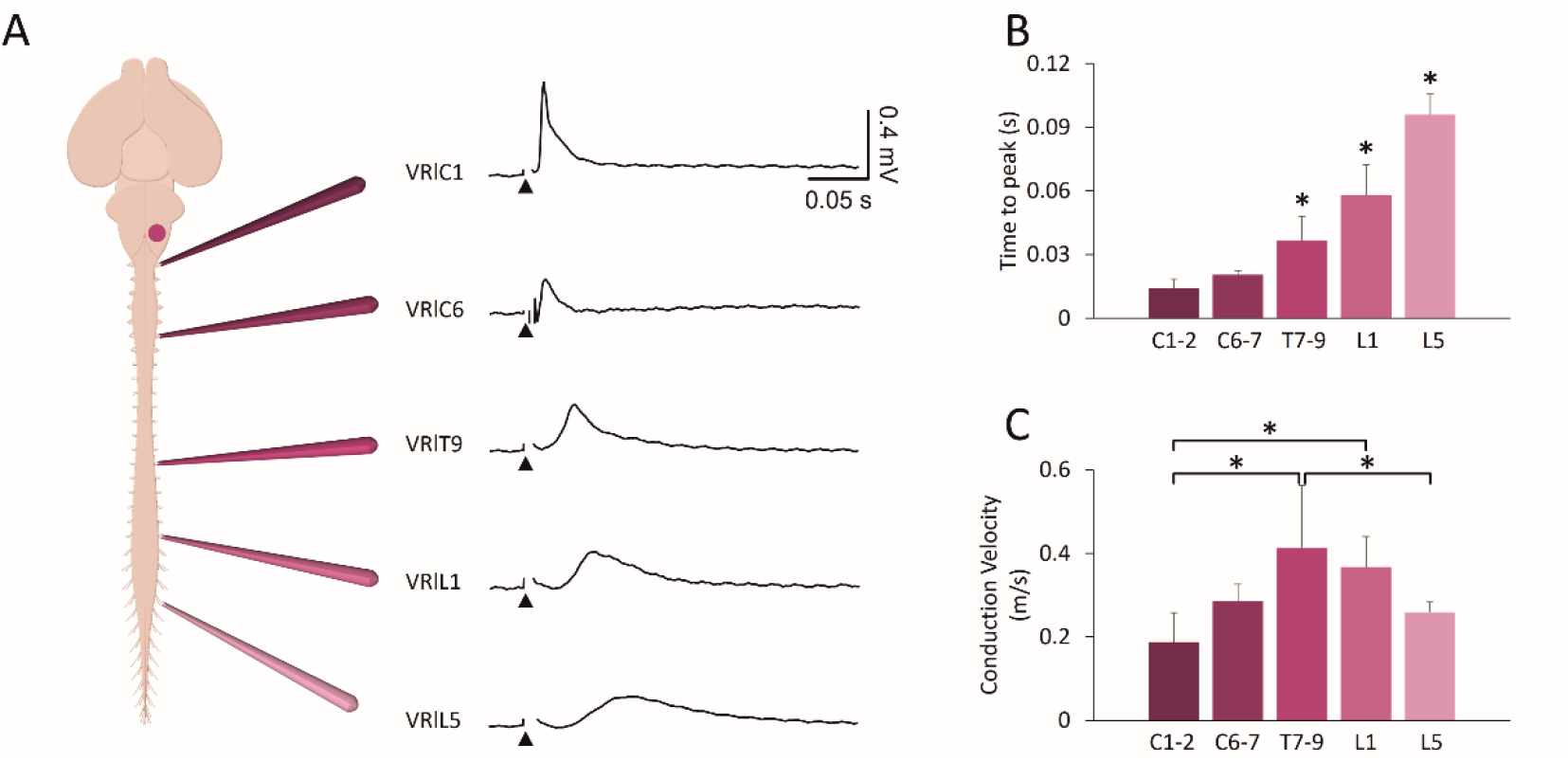
Electrical stimulation of the brainstem reveals different conduction velocities along the cord. (A) Single electrical pulses (intensity = 2 mA, duration = 1 ms) applied to the left pyramids of the brainstem (site H) evokes simultaneous responses from cervical (C1, C6), thoracic (T9), and lumbar (L1, L5) VR on the left side of the cord. Average traces arise from 15 superimposed sweeps. (B) Histogram of the time to peak of responses shows that the slowest response comes from the most caudal segments (*P < 0.001). (C) Histogram for conduction velocity of pulses follow a distinct trend (*P = 0.001).

Data collected from 5 experiments demonstrates that potentials recorded from thoracolumbar segments appear later than cervically-evoked responses (in Fig. 4B; P < 0.001; ANOVA). Pulse conduction velocity based on the actual distance between each pair of stimulating and recording sites revealed that the input descending to thoracic segments is the fastest, and input to lumbar segments is faster than the cervical one (Fig. 4C; P = 0.001; ANOVA; Table 3).

**Table 3.**
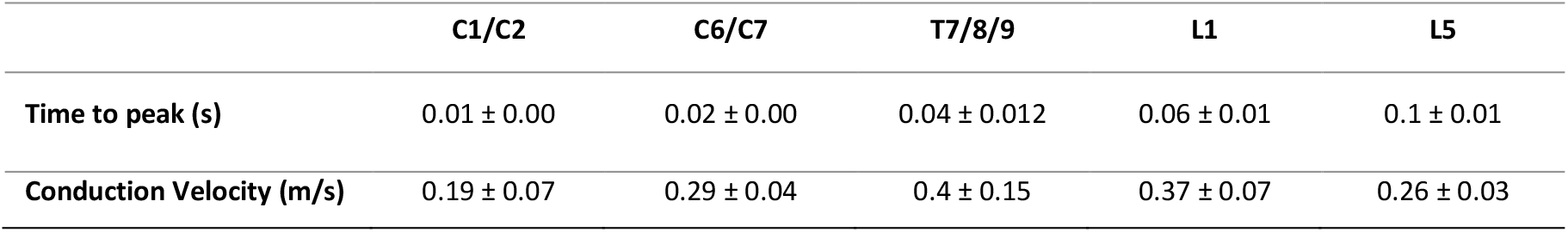
Time to peak and conduction velocity of spinal motor responses evoked by electrical stimulation of left ventrolateral medulla.

The different conduction velocity of descending input elicited by brainstem stimulation suggests that brainstem stimulation enrolls a propriospinal network with distinct synaptic relays.

### Electrical stimulation of caudal dorsal roots evokes ascending input along the cord

After describing the conduction of descending input evoked by brainstem stimulation through the caudal segments of the whole CNS in vitro, we sought to properly characterize the transit of ascending input. For the purpose, electrical pulses were delivered to caudal afferents while monitoring motor responses from rostral segments. In the sample experiment schematized by the cartoon in Fig. 5A, brief pulses (duration = 0.1 ms) were serially applied (0.03 Hz) to a lower lumbar (L5) DR, simultaneously recording responses from homologous and contralateral VRs and from a rostral cervical segment (C2). Input/output stimulation was supplied at augmenting intensities, expressed as multiples of the Thr calculated from the homologous lumbar VR (1 x, 1.5 x, 2 x, 3 x, 5 x Thr), evoking increasingly larger and faster potentials from all spinal VRs. At all strengths of stimulation, motor evoked responses were higher from the homologous VR (Fig. 5B) than from the contralateral L5 (Fig. 5C, note the lower scale on y axis). Motor evoked responses were also induced from upper cervical segments, albeit with lower and delayed potentials (Fig. 5D).

**Figure 5.**
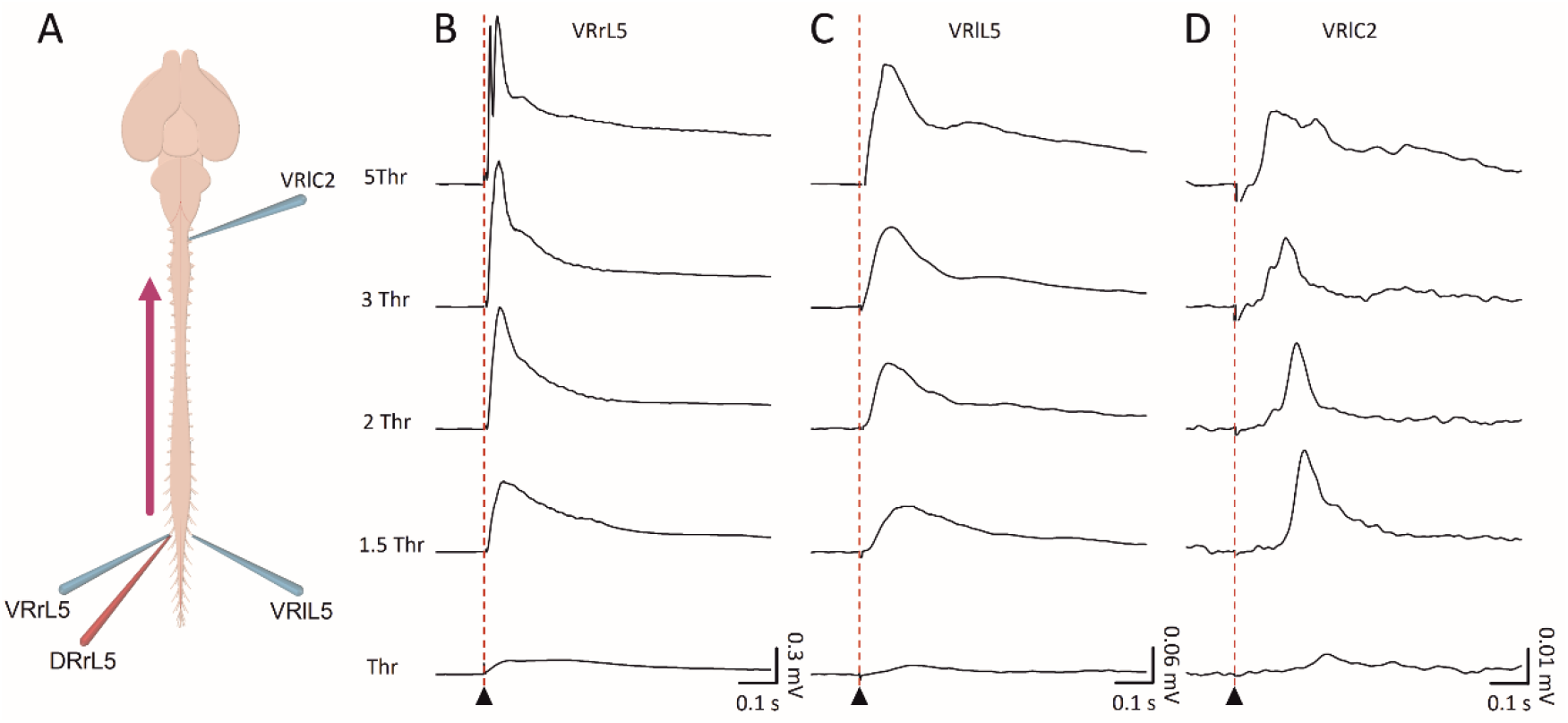
Input output caudal stimulation elicit evoked motor responses along the cord. Electrical pulses (intensity = 0.02-0.1 ms, duration = 0.1 ms) delivered to lumbar afferents of the L5 segment evoke responses from the homologous VRL5 (B), contralateral VRL5 (C) and contralateral higher cervical VRC2 (D). By increasing the strength of stimulation, responses appear sooner and appear higher. Average traces have been pooled from 15 superimposed sweeps.

Input/output average values for each VR are reported in Table 4 as for amplitude and time to peak. Increases in stimulation strengths correspond to higher average peaks of lumbar DRVRPs, while cervical responses remain less affected. The average time to peak of all reflexes was slightly reduced at the first step of increasing stimulation (1 x Thr Vs 1.5 x Thr), while remained unchanged for stronger pulses. Conduction of ascending input along the cord is maintained in the isolated CNS preparation, with cranial VRs far from the stimulating segment showing smaller responses with a late onset. However, no responses from the surface of dorsal motor cortices were acquired in correspondence to dorsal stimulation, even at higher strengths and durations of pulses (data not shown), reminiscent of the above-mentioned inability to elicit spinal responses through electrical stimulation of the motor cortex.

**Table 4.**
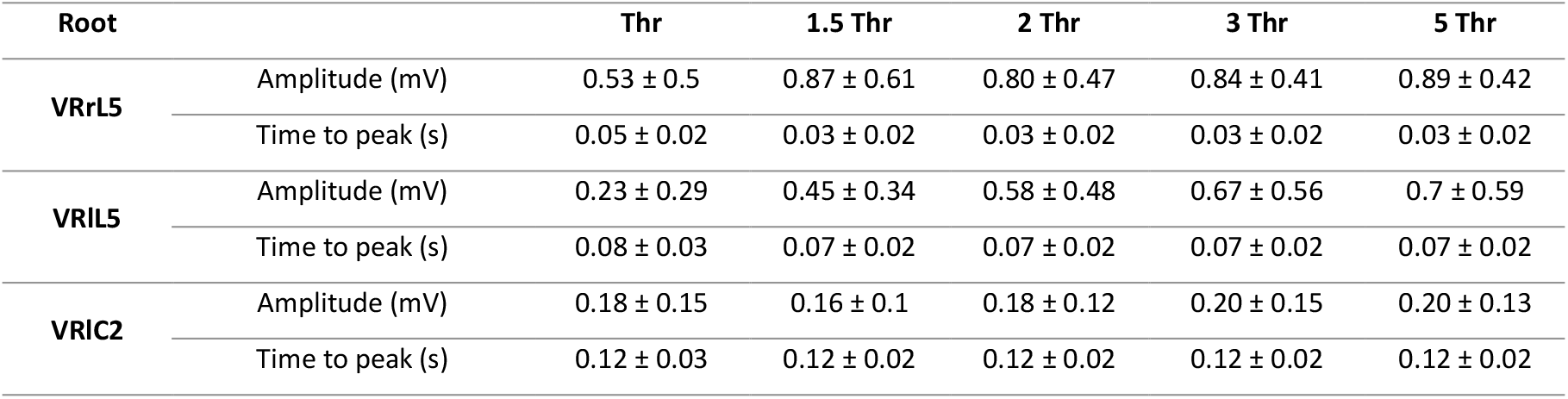
Input-output experiments for electrical stimulation of DRrL5.

### Fictive locomotor patterns are induced by trains of electrical pulses applied to both caudal afferents and ventrolateral medulla

Fictive Locomotion (Kiehn, 2006), consisting in rhythmic electrical oscillations alternating at segmental level between the two sides of the cord and, on the same side, among flexor and extensor motor pools, is a distinctive feature of the activation of neuronal circuits for locomotion in reduced spinal cord preparations (Cazalets et al., 1992). To ascertain whether the recruitment of locomotor spinal networks in vitro is preserved even in the presence of supra-pontine structures, we applied the canonical pattern of electrical stimulation (stereotyped trains of rectangular and brief pulses at 2 Hz) to a caudal afferent of the entire isolated CNS. Sample traces in Fig 6A were taken after about one and half hours from the beginning of the surgical procedures required for the isolation of the entire CNS, and show a cumulative depolarization evoked from four lumbar VRs when a train of rectangular pulses (duration = 0.1 ms) at 2 Hz (blue bar) lasting 80 s was applied to DRrL6. At the top of the cumulative depolarization appeared an epoch of 34 rhythmic discharges in VRlL5, with a mean frequency of 0.45 Hz, that alternated between the flexor-related VRL1 and the extensor-related VRL5 on the right side of the cord, and between bilateral VRL5s. In the same preparation, after transecting the spinal cord at T3/T4 level (Fig. 6B, green bar), the same stimulating protocol elicited a similar episode of FL provided of 28 rhythmic oscillations with a mean frequency of 0.49 Hz in VRlL5. CCF analysis suggests a stronger phase coupling in the intact preparation (CCF_lL2-rL2_ = -0.78, CCF_lL2-lL5_= -0.85) than after midthoracic transection (CCF_lL2-rL2_ = -0.65, CCF_lL2-lL5_ = -0.66).

**Figure 6.**
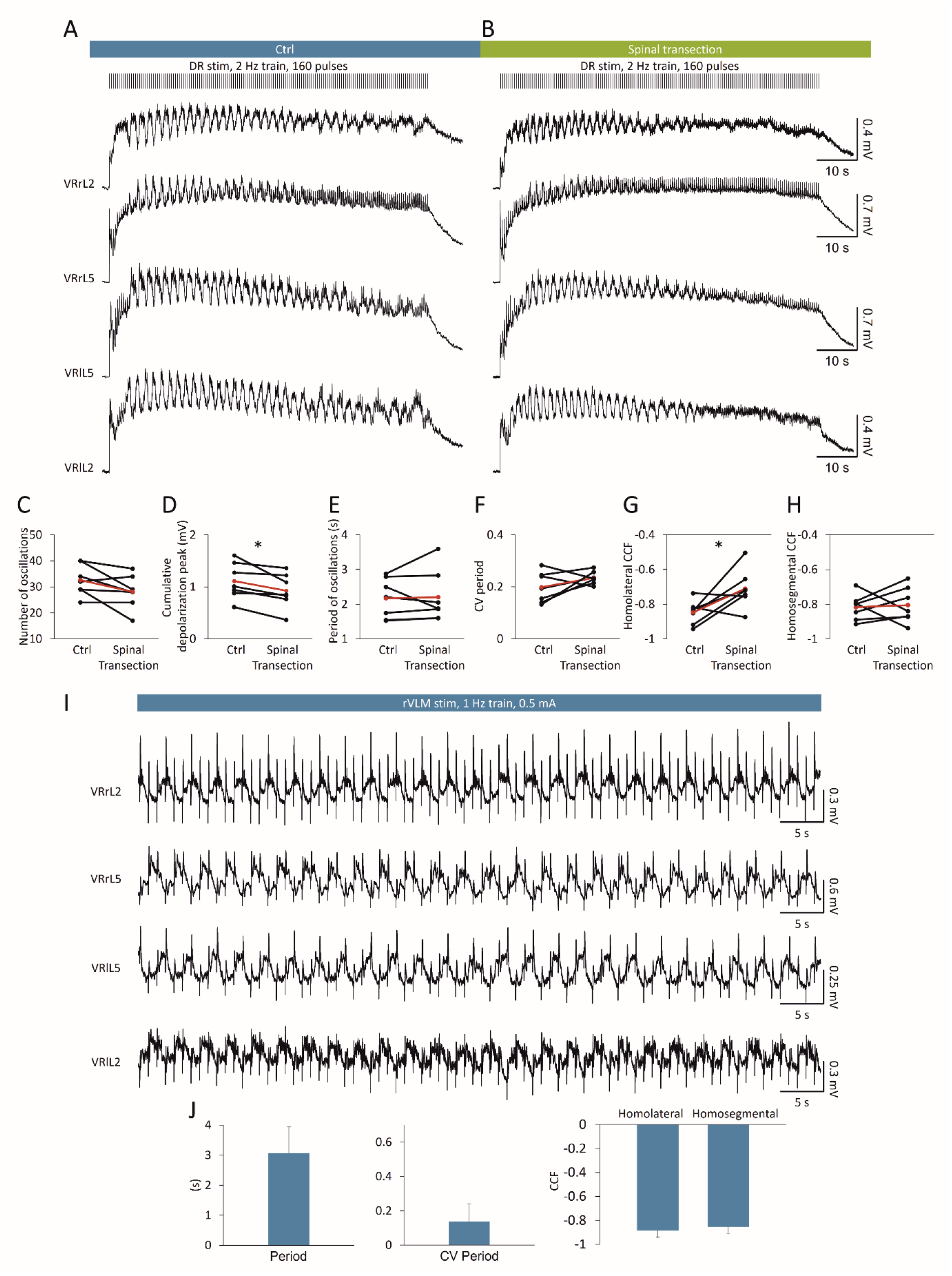
Fictive locomotor patterns are elicited by repetitive electrical stimulation applied either to a lumbar DR or to the ventrolateral medulla. An epoch of FL is induced by a train of pulses (160 stimuli, 2 Hz, intensity = 22.5 µA, pulse duration = 0.1 ms) applied to DRrL6 both in control (A) and after a midthoracic transection (B). VR oscillations appear double alternated between homolateral L2 and L5 segments, and between homosegmental left and right motor pools. Pooled data from seven experiments indicate that, after a midthoracic transection, there are significant differences both in cumulative depolarization peak amplitude (D, *P = 0.023) and in homolateral CCF (cross correlation function; G, *P = 0.038), with an unchanged number of oscillations (C, *P = 0. 075), period of oscillations (E, *P = 0.843), CV period (F, *P = 0.189) and homosegmental CCF (H, *P = 0.797). Red dots indicate the mean values in each graph. (I) Fictive locomotion is stably evoked by the continuous repetitive stimulation (1 Hz) of the right ventrolateral medulla (rVLM, “H” site; intensity = 0.5 mA, pulse duration = 1ms). (J) Bars describe period, CV period, homolateral and homosegmental CCFs of fictive locomotion oscillations as an average of four experiments.

In seven preparations, FL episodes were evoked by a train of 160 pulses at 2 Hz (intensity = 7.5-37.5 µA, 1.5-3.5 Thr; pulse duration = 0.1 ms) applied to lumbosacral afferents (DRrL6, DRrS1). Episodes of FL were the same before and after a midthoracic transection, as for mean number of locomotor-like oscillations (Fig. 6C; P = 0.075, paired t-test), mean period of oscillations (Fig. 6E; period: P = 0.843, paired t-test), and mean variability of cycles (Fig. 6F; CV period: P = 0.189, paired t-test), left and right alternation (Fig. 6H; homosegmental CCF: P = 0.797, paired t-test).

However, total duration of oscillations (67.50 ± 15.82 s vs. 57.61 ± 13.37 s; P = 0.027, paired t-test), mean cumulative depolarization peak (Fig. 6D; P = 0.023, paired t-test) and flexor-extensor alternation (Fig. 6G; P = 0.038, paired t-test) are significantly impaired by a spinal transection. As opposed to transient epochs of FL patterns evoked in the isolated spinal cord, which spontaneously decayed despite the continuous presence of stimulation (Dose et al., 2014), more stable locomotor-like oscillations were obtained in the brainstem spinal cord preparation in response to a train of electrical pulses applied to the ventrolateral medulla (VLM; Zaporozhets et al., 2004). To verify whether electrical stimulation of the VLM induces stable FL patterns in the entire CNS preparation, a train (intensity = 0.5 mA, duration = 1 ms, frequency = 1 Hz) of pulses was continuously applied to the VLM for a total duration of 12 min. In response to stimulation, stable discharges appeared at 0.33 Hz alternating between homolateral extensor and flexor output (CCF = -0.96) and homosegmental left and right ventral roots (CCF = -0.91; Fig. 6I). The same experiment was repeated in four isolated preparations of the entire CNS, where FL patterns stably appeared for up to 12 mins of continuous stimulations (intensity = 0.5 - 4.5 mA, duration = 1 - 5 ms, frequency = 1 - 2 Hz). Locomotor-like events were characterized by stable discharges (CV period value, Fig. 6J) with a frequency of 0.35 ± 0.09 Hz (mean cycle period, Fig. 6J) and double alternating between pairs of VRs (homolateral and homosegmental CCFs, Fig. 6J).

Collectively, FL patterns are evoked in the whole CNS in vitro by trains of electrical pulses applied either to lumbosacral afferents or to the VLM, proving this isolated preparation as a suitable model to study spinal circuits for locomotion in a more intact environment.

### Supra-pontine structures modulate neuronal networks for respiration

Supraspinal structures control several spinal motor output and essential rhythmic behaviors. To better characterize the impact of higher rostral centers on the neuronal pathways involved in respiration, a spontaneous respiratory rhythm was initially derived from upper cervical segments in the whole CNS in vitro and then after two serial surgical transections, first for the precollicular decerebration and afterward for the ablation of pons (Fig. 7A). The raster plot traces the dynamics of respiratory events throughout the entire experimental protocol (2.5 hrs) in a sample experiment (Fig. 7A). The stable respiratory rhythm frequency (green field) was not affected by a precollicular decerebration (yellow field), while it sped up after the following pontobulbar transection (red field). Average bursts were obtained by superimposing the events in the last 5 min of each phase of the experiment in A (Fig. 7B). Single respiratory bursts in the intact CNS (1.88 s; left) shortened after precollicular decerebration (1.24 s; middle) and remained short after pontobulbar transection (1.03 s; right).

**Figure 7.**
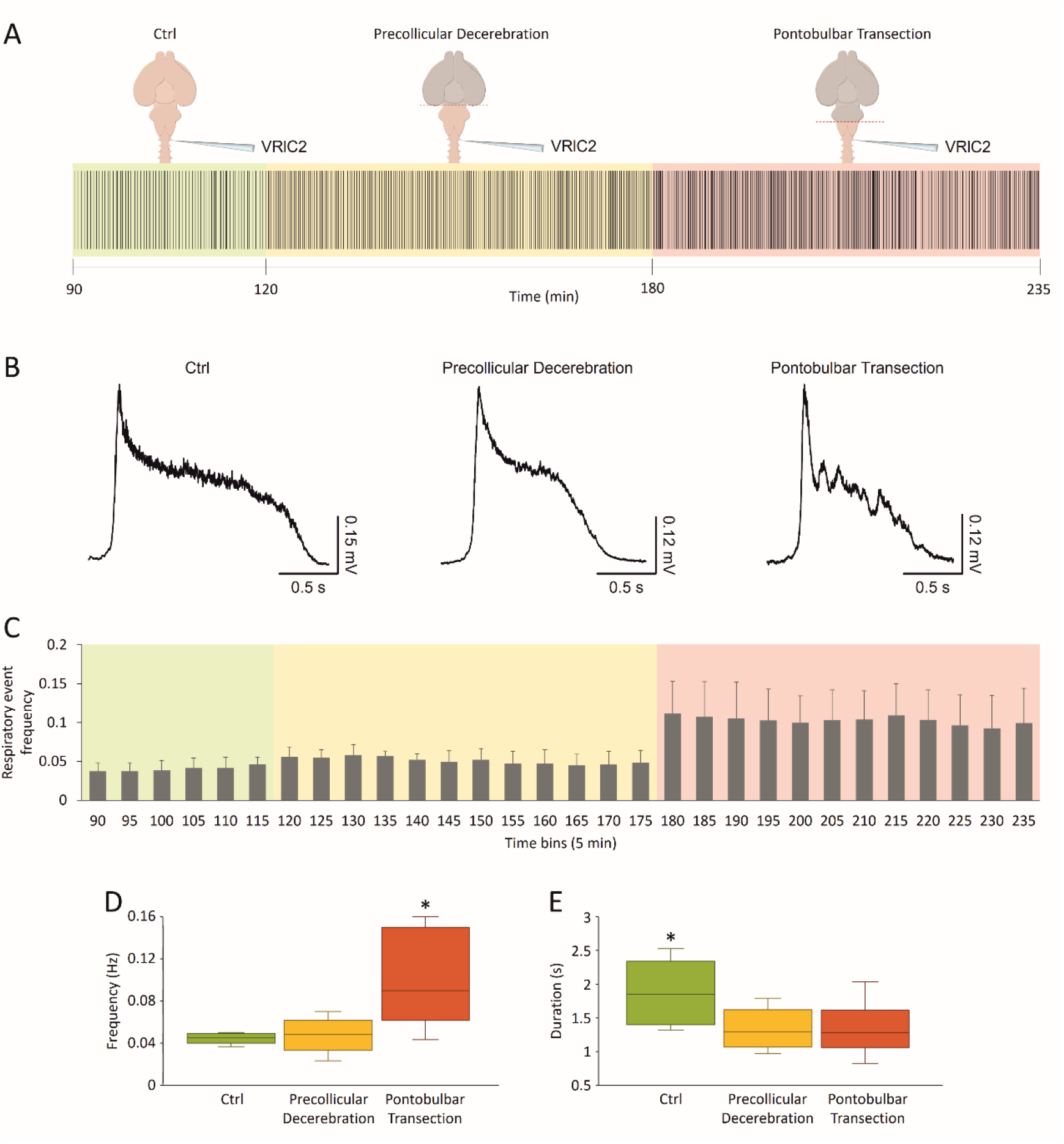
Fictive respiration is modulated by supra-pontine and pontomedullary structures. (A) Raster plot of continuous fictive respiration recorded from VRlC2 for an entire experiment. Time is calibrated at the onset of anesthesia. Fictive respiration in control (green pale background) remained stable after a precollicular decerebration (yellow pale background) and eventually moves faster after the following pontobulbar transection (red pale background). (B) Average bursts from the same experiment in A are reported for the last five minutes of recordings in each experimental slot. Single events become shorter after precollicular decerebration, without any changes after the following pontobulbar transection. (C) An average time course from 8 experiments traces rhythm frequency in 5 min bins for the entire duration of experiments. The last five minutes of each experimental phase in C is used for statistical comparison of rhythm frequency (D, *P = 0.002) and burst duration (E, *P = 0.016).

Time course for the mean frequency from eight experiments is reported for 5 min bins in Fig. 7C. The frequency of the respiratory rhythm increased after the pontobulbar transection (Fig. 7D; P = 0.002; repeated measure ANOVA followed by Tukey-Kramer multiple comparisons test), while burst duration was already reduced after precollicular decerebration (Fig. 7E; P = 0.016, repeated measure ANOVA followed by Tukey-Kramer multiple comparisons test). Collectively, data indicates that supra-pontine structures affect distinct features of the fictive respiratory rhythm, supporting the adoption of the whole CNS in vitro preparation to clarifying the rostral modulation of brainstem networks.

### Supra-pontine structures modulate local lumbar circuitry

We thus explored whether the presence of supra-pontine structures in the CNS preparation affects spinal motor networks in the lumbar cord. To characterize the state of excitability of spinal motor networks, we used dorsally evoked potentials derived from VRs in response to single electrical pulses supplied to dorsal afferents (Lev-Tov and Pinco, 1992). Interestingly, afferent dorsal pulses also recruit a specific dorsal network along the cord (Taccola and Nistri, 2005) that is involved in the presynaptic inhibition of input coming from the periphery (Rudomin, 2009).

As shown in the sample mean traces reported in Fig. 8A and B, electrical stimulation (6-60 µA 2-3.3 Thr, 100 µs) of a DRlL5 in the isolated CNS elicits an early sharp peak from VRs coming essentially from an oligosynaptic pathway in the local microcircuitry, and a following long potential corresponding to the activation of a larger number of interneurons. A sharper potential is induced also from DRlL2 (Fig. 8C), following antidromic conduction of the primary afferent depolarization elicited by DRlL5 stimulation. Sample average traces in Figure 8A-C were acquired before and after a precollicular transection, indicating that decerebration causes a faster decay of all responses and a 23% reduction in the peak of responses from VRrL2 (Fig. 8B).

**Figure 8.**
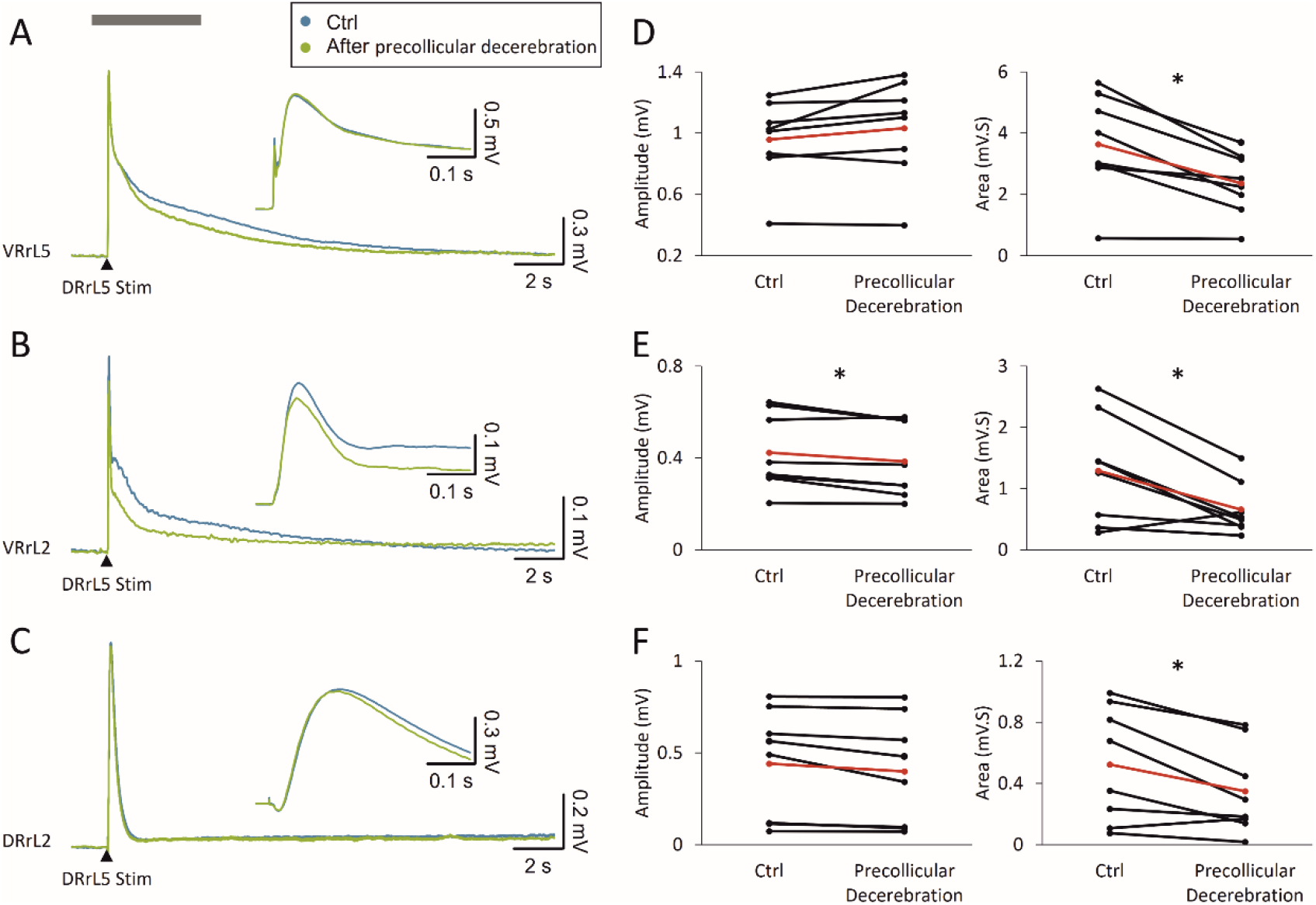
Local lumbar networks involved in reflex responses are modulated by supra-pontine structures. Superimposed average traces show that electrical pulses applied to a DRlL5 (intensity = 45 µA, pulse duration = 0.1 ms) induce both VR and DR potentials from VRrL5 (A), VRrL2 (B) and DRrL2 (C). After acquiring evoked responses in control (blue traces), the same stimulating protocol is repeated after precollicular decerebration (green traces). Inserts show the magnified onset of response to better appreciate any changes in peak amplitude after decerebration. (D-F) For each recording site, pairs of responses before and after decerebration are quantified for amplitude and area. Red dots correspond to the mean value in each graph (D, *P = 0.003; E, amplitude *P = 0.016, area *P = 0.039; F, *P = 0.016).

Reflexes were acquired from both the intact CNS preparation (blue lines) and after precollicular decerebration on the same sample (green lines). As summarized in the plots of Fig 8D-F, decerebration does not affect peak amplitude from VRrL5 (P = 0.103, paired, t-test; n = 8) nor DRrL2 (P = 0.055, paired, t-test; n = 8), while peak amplitude is reduced from VRrL2 (P = 0.016 paired, t-test; n = 8).

Similarly, the area of all responses is depressed after precollicular transection (Area_VRrL2_ P = 0.039, Wilcoxon matched-pairs signed-ranks test; Area_VRrL5_ paired t-test, P = 0.003; Area_DRrL2_ P = 0.016 paired t-test; n = 8).

These observations demonstrate that the lack of supra-pontine structures deprives lumbar circuits of some modulatory influences, spurring the adoption of the whole in vitro CNS preparation whenever interested in exploring supraspinal influences on spinal microcircuits.

### The isolated CNS with legs attached expresses a stable spontaneous fictive respiration modulated by supra-pontine structures

Isolated preparations of brainstem and spinal cord with hindlimbs kept intact and connected to the spine have been used to explore both the functional coupling between networks for respiration and locomotion (Giraudin et al., 2012), as well as the modulatory influence on spinal circuits played by the afferent feedback elicited by passive exercise (Dingu et al., 2018). However, to explore whether supra-pontine structures contribute in integrating the afferent input elicited by passive leg movement, the definition of a more intact in vitro preparation became compelling. To this purpose, we set a semi-intact preparation of the whole CNS with intact hindlimbs connected to the spine (Fig. 9A), which expresses a stable spontaneous respiratory rhythm for over 4 hours when surgical procedures for tissue isolation are fast (≤40 min) and performed on younger animals (≤2 day old), summarized in the scatter plot of Fig. 9B from 66 preparations. In Fig. 9C, the Raster plot describes a sample respiratory rhythm from a two-day old neonatal preparation, which is progressively slowed down by precollicular decerebration (Fig. 9C, yellow field) and then speeded up by the following pontobulbar transection (Fig. 9C, red field). Original traces at steady state illustrate the regular bursting at 0.08 Hz in the intact preparation (Fig. 9D), which slowed down to 0.06 Hz after precollicular transection (Fig. 9E) and then accelerated to 0.10 Hz following ablation of the pons (Fig. 9D). In the intact preparation, the average single burst lasted 1.63 s (Fig. 9D_1_) and was reduced to 1.43 s after ablation of supra-pontine structures (Fig. 9E_1_), while pontobulbar transection broadened durst duration to 1.84 s (Fig. 9F_1_). Pooled data from many experiments confirms the significant reduction of bursting frequency after precollicular decerebration (from 0.07 ± 0.02 Hz to 0.05 ± 0.02 Hz; p = 0.036; One way ANOVA followed by all pairwise multiple comparisons with Fisher LSD Method; n = 9) and its subsequent recovery after pontobulbar transection (0.07 ± 0.02 Hz; p = 0.036; One way ANOVA followed by all pairwise multiple comparisons with Fisher LSD Method; n = 6). Similarly, burst duration was reduced by decerebration (from 1.72 ± 0.46 s to 1.22 ± 0.43 s; p = 0.003; One way ANOVA on ranks followed by all pairwise multiple comparisons with Dunn’s Method; n = 11) and then broadened again after the following pontobulbar transection (1.68 ± 0.25; p = 0.003; One way ANOVA on ranks followed by all pairwise multiple comparisons with Dunn’s Method; n = 8). In summary, hindlimbs kept attached to the isolated CNS do not affect the expression of the spontaneous respiratory rhythm, nor its modulation provided by supra-pontine structures. Thus, the whole CNS with legs attached represents an original setting to explore how respiration is tuned by sensory input reaching the brain from the periphery.

**Figure 9.**
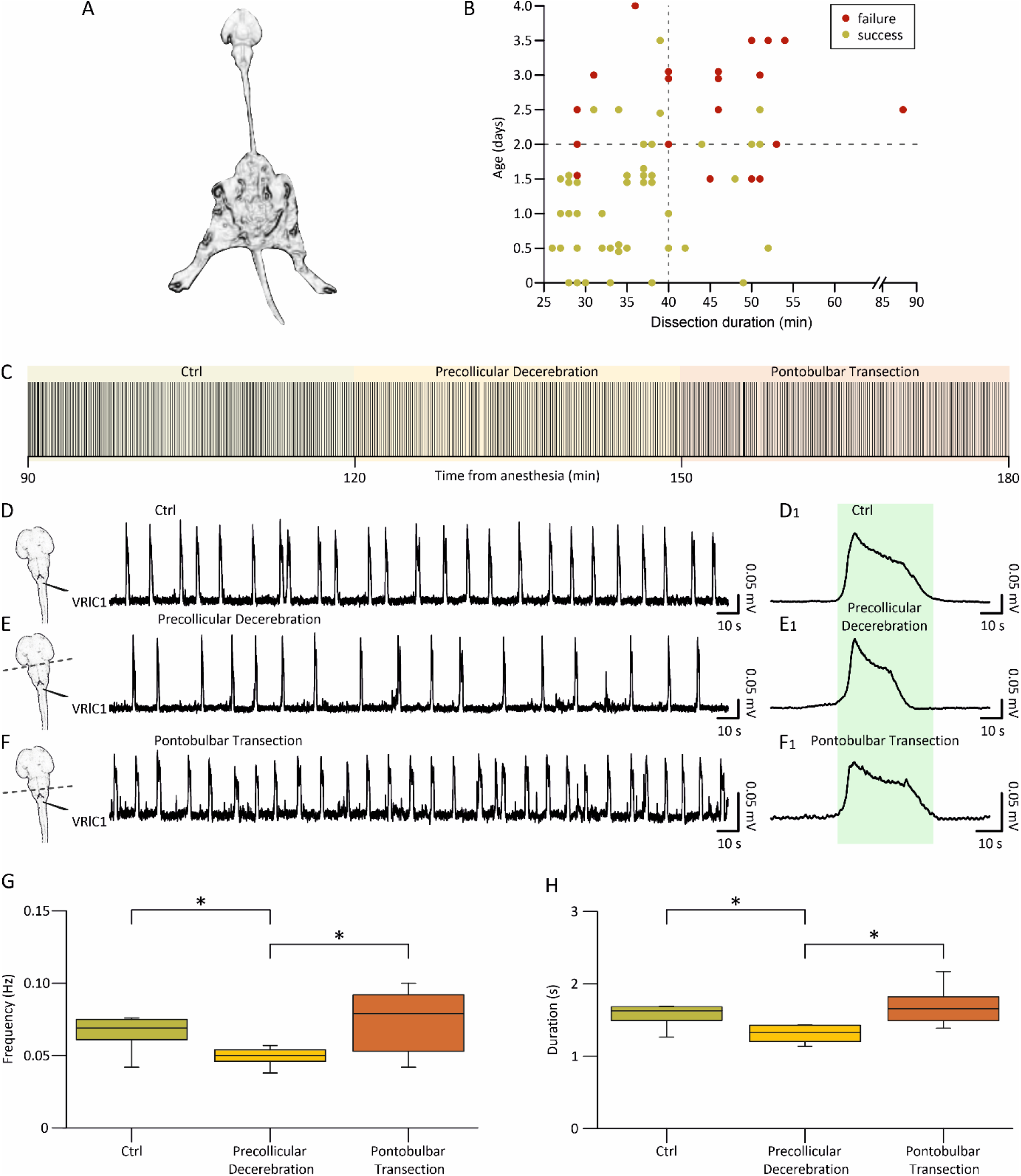
In vitro preparation of the entire CNS with hindlimbs attached expresses a stable fictive respiration that is affected by precollicular and pontomedullary serial transections. (A) Picture of the original in vitro preparation from a neonatal rat (P2), comprising the whole central nervous system with hindlimbs attached. Lower extremities and tail were left intact, along with ventral roots (VRs) and dorsal roots (DRs) below Th13 segment. (B) Plot depicts each preparation as a single dot (66 preparations) describing age of the animal (Y axis) and length of surgical dissection (X axis). Dots are colored in green (success) or red (failure) based on the presence or absence of a stable respiratory rhythm after at least 4 hours from the induction of anesthesia at the beginning of tissue isolation. Vertical dotted gray line at x = 40 min and horizontal dotted gray line at y = 2 days, define a bottom left quadrant where the probability of having preparations with long-lasting breathing is highly consistent (44 successes out of 47 preparations). (C) Raster plot representing consecutive respiratory bursts continuously recorded from a cervical VR (lC1) in the intact CNS (ctrl; 30 min; green shadow field) and after serial precollicular (30 min; orange shadow field) and pontobulbar transections (30 min; red shadow field). Albeit the expected variability, rhythm frequency is reduced after decerebration, while it is increased by the subsequent pontomedullary transection. For the same experiments reported in C, three trace segments from VRlC1 are taken at steady state in intact settings (D) and during the progressive reductions of the intact CNS preparation (E, F; see left cartoons). Burst frequency in the entire CNS (top trace) is slowed down by decerebration (middle trace) and then eventually speeded up after pontomedullary transection (bottom trace). (D_1_, E_1_, F_1_) Average bursts calculated by superimposing single events are reported on the right, demonstrating that decerebration reduces burst duration, which eventually recovers after further tissue ablation. Differences in burst duration are highlighted by the green shaded field corresponding to the burst duration calculated in control. Effect of rostral structures ablation on the frequency of respiration is summarized by whisker plots from pooled data in G (n = 9), showing significant changes in the pace of rhythm from the intact CNS (green box, ctrl, n = 9) following precollicular decerebration (yellow box, precollicular transection, n = 9) and then pontomedullary transection (orange box, pontobulbar decerebration, n = 6; *P = 0.036). H. Whisker plots describing changes in burst duration from the intact CNS (green box, n = 11) following serial transections (yellow box, precollicular transection, n = 11; orange box, pontobulbar decerebration, n = 8; *P = 0.003). Note that burst amplitude has not been evaluated since nerves were released from electrodes before each surgical ablation to avoid root damage.

## Acknowledgments

GT is also grateful to Mrs. Elisa Ius for her excellent assistance in preparing the manuscript. We are grateful to Dr. Graciela Mazzone for helpful data discussion.

## Abbreviations

ANOVA: analysis of variance
CCF: cross correlation function
C: cervical
CNS: central nervous system
DR: dorsal root
DRDRP: dorsal root dorsal root potential
DRVRP: dorsal root ventral root potential
l: left
L: lumbar
P: postnatal
r: right
Thr: threshold
T: thoracic
VR: ventral root

## Notes

### Competing Interest Statement

The authors have declared no competing interest.

